# Validation of predicted anonymous proteins simply using Fisher’s exact test

**DOI:** 10.1101/2021.04.28.441814

**Authors:** Jean-Michel Claverie, Sébastien Santini

## Abstract

**Motivation:** Genomes sequencing has become the primary (and often the sole) experimental method to characterize newly discovered organisms, in particular from the microbial world (bacteria, archaea, viruses). This generates an ever increasing number of predicted proteins the existence of which is unwarranted, in particular among those without homolog in model organisms. As a last resort, the computation of the selection pressure from pairwise alignments of the corresponding “Open Reading Frames” (ORFs) can be used to validate their existences. However, this approach is error-prone, as not usually associated with a significance test.

**Results:** We introduce the use of the straightforward Fisher’s exact test as a post processing of the results provided by the popular CODEML sequence comparison software. The respective rates of nucleotide changes at the non-synonymous vs. synonymous position (as determined by CODEML), are turned into entries into a 2×2 contingency table, the probability of which is computed under the Null hypothesis that they should not behave differently if the ORFs do not encode actual proteins. Using the genome sequences of two recently isolated giant viruses, we show that strong negative selection pressures do not always provide a solid argument in favor of the existence of proteins.

## 1 Introduction

Since the first two bacterial genomes were sequenced 25 years ago (Fleischmann *et al*., 1995; Fraser *et al*., 1995), partial and whole genome sequencing have become the method of choice in identifying and characterizing new microorganisms (bacteria, archaea, unicellular eukaryotes, viruses), revealing the stupendous extent of their diversity. By their simplicity of use and low cost, the most recent 3rd generation sequencing platforms (Goodwin *et al*., 2016) have made microbial genomics accessible to non-specialists (MacLean *et al*., 2009; Gwinn *et al*., 2019), while a few large centers are fully taking advantage of their huge throughput to run biodiversity exploration projects of ever increasing dimensions (Lewin *et al.* 2018; Sunagawa *et al*., 2020; Chen *et al.* 2021). The most recurring (and unexpected) lesson emerging from the analyses of these enormous datasets is that overall morphological and phylogenetic similarities, as well as similar metabolisms and lifestyles, could hide large differences in gene contents and encoded proteomes. Within microorganisms belonging to a given clade, such as eukaryotic classes, bacterial genera, or virus families, genomes are found to encode a subset of « core » proteins (i.e. with homologues in all members) together with proteins unevenly distributed, some of which only present in a single species or strain. This dichotomy is best documented for bacteria and viruses for which core genes might only represent a small proportion of the pangenome, i.e. of all the genes occurring at least once among all clade members (Land *et al*., 2015, Claverie & Abergel, 2018). While there is little doubt that genes encoding proteins with homologs in multiple divergent members of a clade are real, the level of certainty is much lower when they only occur once, or within very close clade members where the corresponding Open Reading Frames (ORF) may occur by chance. Given the A+T richness of STOP codons (TAA, TAG, TGA), random ORFs are also statistically expected to occur at higher frequency in high G+C content genomes, increasing the risk of protein overprediction (Legendre *et al*., 2019). The uncertainty further increases when the predicted proteins are short (typically less than 100 residues), or do not exhibit neither a functional motif nor a significant sequence similarity in the reference databases (Sayers *et al*., 2021). Such cases, referred to as “ORFans” represent a large proportion of predicted microbial proteomes (Entwistle *et al*., 2019) in particular for large viruses (Philippe *et al*., 2013; Gallot-Lavallée *et al*., 2017; Legendre *et al*., 2018; Abergel & Claverie, 2020; Boratto *et al*., 2020). The validation of ORFans is important to document the intriguing evolutionary process of *de novo* gene creations from non-coding regions in prokaryotes, eukaryotes, and their viruses (Legendre et al., 2018; McLysaght & Guerzoni, 2015; Schlötterer, 2015; Schmitz & Bornberg-Bauer, 2017; Van Oss & Carvunis, 2019).

If the experimental validation of predicted proteins through mass-spectrometry has become easier, the technique remains inaccessible to many of the laboratories generating genomic data. It also requires the corresponding microorganisms to be isolated and cultivated, thus disregarding the increasing number of metagenomics assemblies (Benler *et al.* 2021). Furthermore, certain proteins might only be expressed (and experimentally detectable) at specific time in the life cycle of an organisms, in certain environmental conditions, or in specific organs. Thus, our capacity of experimentally demonstrating the actual existence of predicted proteins has fallen much behind the overwhelming production of genomic data. To overcome this difficulty it has become customary to compute the selection pressure, i.e. the ratio of the synonymous vs. non-synonymous mutation rates as a way to validate bioinformatic protein predictions (e.g.: Doutre *et al*., 2014: González *et al*., 2016); Prabh & Rödelsperger, 2016); Christo-Foroux *et al*., 2020).

The concept/calculation of the selection pressure is based on the fact that proteins are made for a purpose and that their functions, directly derived from their amino-acid sequences, tend to be conserved throughout evolution. Accordingly, we expect that the non-synonymous positions of their coding regions will vary much less rapidly than the synonymous ones, the changes of which have lesser consequence on the organism’s fitness.

The concept of selection pressure, was most widely disseminated via the CODEML program of the PAML package for phylogenetic analysis (Yang, 2007; Yang, 2014; Jeffares, 2015). The computation requires the disposal of at least two homologous ORF sequences, and involves five straightforward steps: 1) from the comparison with the associated aminoacid sequence, each position in the ORF nucleotide sequence is classified as synonymous or non-synonymous in reference to the degeneracy of the genetic code. Their respective total numbers are denoted N_S_ and N_NS_; 2) the two homologous amino-acid sequences are optimally aligned, then codon-wise converted into a nucleotide sequence alignment; 3) the observed nucleotide changes associated to the positions previously mapped as synonymous or non-synonymous are separately counted and are denoted nS and nNS; 4) one then forms the ratios d_N= n_NS__/N_NS_ and d_S_= n_S_/N_S_, separately quantifying the mutation rates at the two different types of positions; 5) finally, one compute the “selection pressure” as the ratio ω= d_N_/d_S_.

The values of ω are intuitively interpreted as follows: ω < 1 will correspond to proteins the (beneficial) functions of which resist amino-acid changes, also said to evolve under negative (purifying) selection. This is by far the most frequent situation. In contrast, ω > 1 correspond to the less frequent cases where changes in protein are positively selected (i.e. adaptive evolution) either to modify or abolish its (detrimental) function. Although conceptually simple, the practical implementation of this analysis comes up against two contradictory constraints. The first is that it must be based on an alignment of impeccable quality, and therefore between two highly similar protein sequences. The second is that the number of substitutions must be sufficiently high, while keeping the probability of multiple substitutions at the same site negligible (which would distort the estimate of d_s_ and d_ns_). To our knowledge, the validity range of the method was never rigorously defined in terms of pair-wise sequence divergence (i.e. acceptable value ranges for NS, NNS, and the dS or dN ratios), although CODEML can compute a likelihood value for a large suite of adaptive evolution models (the grasp of which is beyond the reach of most of occasional users). Fortunately, the use of CODEML remains easily tractable (Jeffares, 2015) if we only wish to compute ω from the pairwise alignment of two homologous ORFs in order to evaluate the quality of *ab initio* protein prediction, as presented in the next section.

## 2 Methods

For actual proteins, the non-synonymous and synonymous positions of the coding regions are expected to diverge at different speeds, thus leading to ω ≠1 in most cases. In contrast, in the case of false protein (ORF) predictions, the bioinformatic distinction made between non-synonymous and synonymous positions becomes irrelevant, and both types of positions are no longer expected to display a different mutational behavior. We then expect ω to remain close to one, within the range of random fluctuations. As the non-synonymous and synonymous positions are two mutually exclusive categories, we can evaluate how much both positions behave differently using Fisher’s exact test in the analysis of the 2×2 contingency table computed from the pairwise alignment of two homologous protein predictions, as follows:

| Position type | # Substituted | # Non-substituted |
| --- | --- | --- |
| Non-synonymous | $n_{NS}$ | $N_{NS} - n_{NS}$ |
| Synonymous | $n_S$ | $N_S - n_S$ |

Where n_NS_ and n_S_ are computed as the products d_N_.N_NS_ and d_S_.N_S_, respectively. These values are directly read from the standard CODEML output, then rounded to the nearest integers to be compatible with Fisher’s test. The probability (p-value) that both position types (synonymous and non-synonymous) behave differently (hence that the ORFS prediction are dubious) can be calculated by any available implementation of Fisher’s test (online or in R, for instance). The pairwise sequence alignments were analyzed using the PAML 4.9j package version for UNIX/Linux with the following relevant options:

noisy=0; verbose=0; runmode=-2 (pairwise); seqtype = 1 (codons); CodonFreq=2; model=1; NSsites=0; icode=0 (universal code); fix_kappa=1; kappa=1; fix_omega=0 (omega to be estimated); omega=0.5.

We apply the above procedure to the evaluation of the whole predicted proteomes of two virus sequenced in our laboratory, constituting the only two known members of the proposed *Molliviridae* giant virus family. The prototype of the family, *Mollivirus sibericum* was isolated from ancient Siberian permafrost (Legendre *et al*., 2015) while the second member, *Mollivirus kamchatka,* was isolated from surface soil in Kamtchatka (Christo-Foroux, *et al*., 2020). Both are “giant” DNA viruses infecting the protozoan *Acanthamoeba.*

A stringent gene annotation of *M. sibericum* was initially performed using transcriptomic data (stranded RNA-seq reads) in addition to the standard protein-coding prediction methods (Legendre *et al*., 2015). *M. kamchatka* proteome prediction (Christo-Foroux, *et al*., 2020) was performed without RNA-seq data but taking into account protein similarity with *M. sibericum*. Gene predictions were further curated using the web-based genomic annotation editing platform Web Apollo (Dunn *et al*., 2019). The selection pressure analysis was performed using CODEML as previously described (Christo-Foroux, *et al*., 2020). Finally, the codon adaptation index (CAI) of both mollivirus predicted proteomes was performed using the CAI tool from the Emboss package (Rice et al., 2000).

## 3 Results

A total of 495 and 480 genes were predicted for *M. sibericum* and *M. kamchatka,* with the encoded proteins ranging from 51 to 2,171 residues and from 57 to 2,176 residues, respectively (Christo-Foroux, *et al*., 2020). While the two isolates are very close to each other, sharing 463 of their predicted proteins as best reciprocal matches (with 92% identical residues on average, using BlastP), they are also very different from all other known organisms with 60% of their predicted proteins lacking a detectable homolog among cellular organisms or previously sequenced viruses (outside of the proposed *Molliviridae* family). These ORFan-rich proteomes constitute an ideal test set for our proposed selection pressure-based validation procedure to distinguish proteins that are actually made from ORFs that might be conserved by chance between evolutionary close viruses. **Figure 1** displays the selection pressure values computed for all pairs of ORFans (panel A) and non-ORFans (panel B). For comparison the exact same pairs of genes were also displayed in association to more traditional parameters such as their length, CAI, and (G+C) %. In all graphs, ORF pairs associated to non-significant Fisher’s test p-values (thus less likely to correspond to actual proteins) are indicated by red dots.

**Fig. 1.**
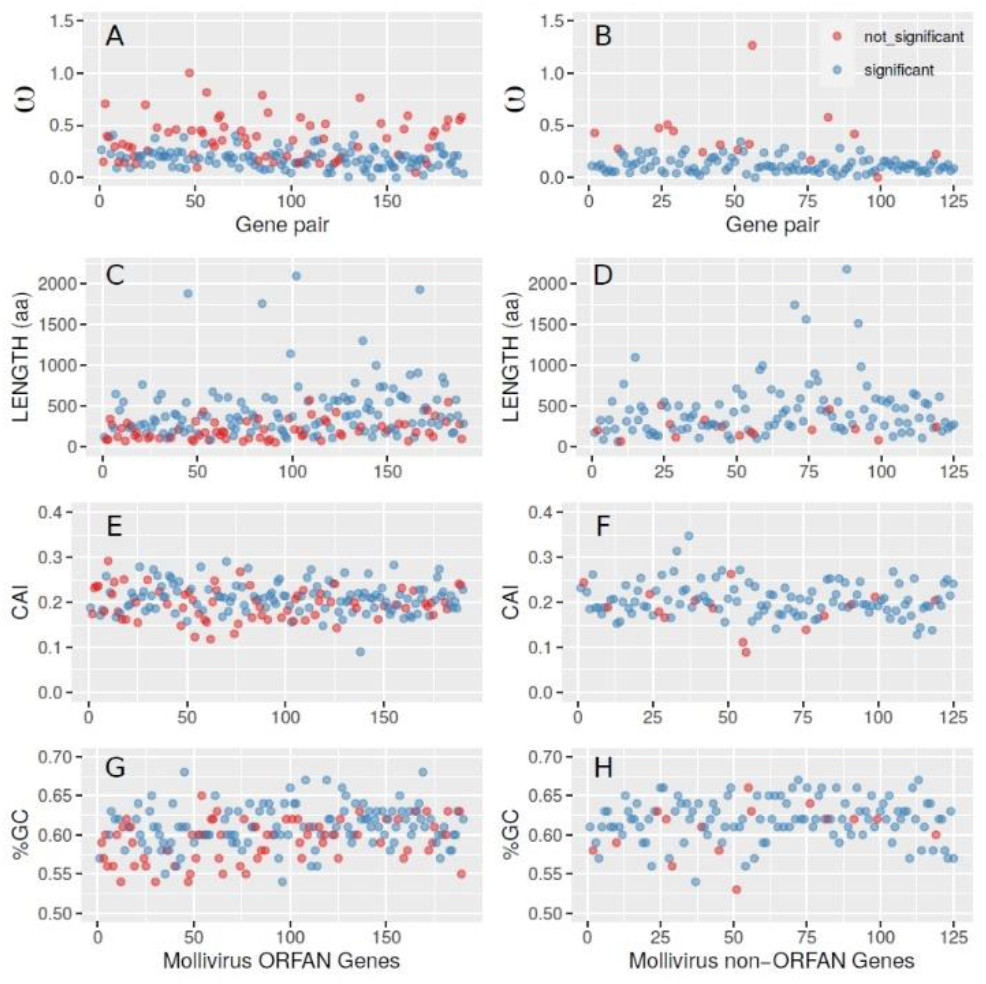
Selection pressure values and other parameters associated to ORFans vs non-ORFans predicted Mollivirus proteins. Each dot corresponds to a pair of orthologous genes, the relative genomic position of which is indicated by the X-axis, separately for each column. The left/right columns correspond to ORFans/non-ORFans, respectively. ORFs associated to ω values not significantly different from 1 are in red (p-value >1/495, allowing for one false positive), others are in blue.

**Table 1** provides some of the numerical values distinguishing the ORFan vs. non ORFans gene populations as well as those associated to ω values non-significantly or significantly different from 1 (i.e. “dubious” *vs.* “confirmed” protein candidates), as displayed in Fig. 1.

**Table 1.** Properties of predicted ORFs associated to ω values significantly (blue) *vs.* non-significantly (red) different from 1 (Note: *K-S: Kolmogorov-Smirnov test).

|  | ORFans |  |  |
| --- | --- | --- | --- |
|  | Blue | Red | K-S* test |
| Total | 122 | 68 | NA |
| $\omega$ | $0.18 \pm 0.09$ | $0.38 \pm 0.2$ | $p < 2. e-11$ |
| Length (aa) | $457 \pm 355$ | $210 \pm 120$ | $p < 3. e-08$ |
| CAI | $0.21 \pm 0.03$ | $0.2 \pm 0.04$ | $p > 0.03$ |
| %G+C | $0.61 \pm 0.03$ | $0.59 \pm 0.03$ | $p > 0.016$ |
|  | Non ORFans |  |  |
| Total | 110 | 15 | NA |
| $\omega$ | $0.12 \pm 0.07$ | $0.4 \pm 0.28$ | $p < 1.1 e-06$ |
| Length (aa) | $444 \pm 338$ | $232 \pm 126$ | $p < 0.01$ |
| CAI | $0.21 \pm 0.04$ | $0.18 \pm 0.05$ | $p > 0.74$ |
| %G+C | $0.62 \pm 0.03$ | $0.61 \pm 0.03$ | $p > 0.87$ |

The most discriminant pattern in Fig. 1 is the larger proportion of red dots in the left columns. ORFans are more frequently associated to ω values non-significantly departing from one (68/190, 36%) than non-ORFans (15/125, 12%). This suggests that more than a third of predicted ORFans might not correspond to actual proteins. Yet, this result also shows that our testing procedure provides confirmations for all the others (64%)(Table 1). The much larger proportion of blue dots for the non-ORFans, confirms that the detection of homologs (even in very close species) is a reliable way to assess the reality of predicted proteins. However, our results indicate that 12% of them might be undergoing pseudogenization, despite appearing to remain under negative selection (ω values < 0.6, Fig. 1B).

The utility of our ω value testing procedure is best illustrated by the combined consultation of Fig. 1 and Table 1. For instance, it is clear that dubious predictions (red ORFs) are in average associated to larger ω value than the blue ones (Table 1, p-values correspond to a Kolmogorov-Smirnov test). However, Fig. 1 (A & B) shows that no unique minimal ω value threshold can be used to cleanly separate the two populations.

Similarly, the predicted protein length distribution are significantly different between the red and blue dots (Table 1). However, if smaller ORF predictions are clearly less reliable, no clear length threshold could separate both distributions (Fig.1 C & D).

Finally, the computations of the CAI (Fig. 1, E & F) or G+C content (Fig. 1, G & H) do not bring in usable information to discriminate between reliable or unreliable protein predictions, given the very similar value distributions of these parameters for the red and blue dot populations.

We started this work by noting that ω is defined as the ratio of two small quantities (dN/dS) themselves computed from a limited number of substitution events (ns, nNS) imposed by the necessity of flawless pairwise sequence alignments. Values of ω are thus highly sensitive to random fluctuations making them unreliable to assess the validity of protein prediction. We showed here that applying the Fischer’s exact test to the standard CODEML output, provides a simple way to improve the reliability and predictive power of pressure selection computations. This procedure might thus constitute a useful addition to the standard genome annotation pipe-line and previously proposed software tools to help identify spurious ORFs (Eberhardt *et al*., 2012; Höps *et al*., 2018). As a side benefit, the use of Fisher’s test automatically filters out pairwise alignments that do not exhibit enough substitutions because they are too similar, or their alignments too short. The only parameter remaining to be fixed is the % of identical residues between orthologous proteins that should be greater than 70% (usually by imposing dS<2) to ensure high quality pairwise alignments and minimize the probability of multiple substitutions at one given site (Jeffares, 2015).

## Acknowledgements

We acknowledge the use of the PACA-Bioinfo platform.

## Funding

The IGS laboratory is partially supported by recurrent funding from CNRS and Aix-Marseille University.

## Software and Data Availability

The PAML 4.9j package used to run CODML is freely available at: http://abacus.gene.ucl.ac.uk/software/paml.html.

The sequence data used here are available at https://www.ncbi.nlm.nih.gov/nuccore/under accession#: NC_027867 and MN812837 for *Mollivirus sibericum* and *Mollivirus kamchatka,* respectively.

## Conflict of Interest

none declared.

## Notes

### Competing Interest Statement

The authors have declared no competing interest.

### Summary of Updates

The procedure is tested and validated on a much bigger data set (two complete viral genomes).

